# Spontaneous Isomerization of Asp387 in Tau is Diagnostic for Alzheimer’s Disease: An Endogenous Indicator of Reduced Autophagic Flux

**DOI:** 10.1101/2021.04.21.440819

**Authors:** Evan E. Hubbard, Lilian Heil, Gennifer E. Merrihew, Jasmeer P. Chhatwal, Martin R. Farlow, Catriona A. McLean, Bernardino Ghetti, Kathy L. Newell, Matthew P. Frosch, Randall J. Bateman, Eric B. Larson, C. Dirk Keene, Thomas J. Montine, Michael MacCoss, Ryan R. Julian

## Abstract

Amino acid isomerization is a spontaneous chemical modification potentially related to the underlying causes of Alzheimer’s disease (AD). We demonstrate that data-independent acquisition mass spectrometry can be used to characterize isomerization in complex protein mixtures. Examination of a large cohort of brain tissue samples revealed a striking relationship between isomerization of tau and AD status. Surprisingly, isomerization was found to be more abundant in both autosomal dominant and sporadic AD samples relative to controls. We hypothesize that lower autophagic flux in AD brains accounts for these results. Additional data, including quantitative analysis of proteins related to autophagy, strongly support this hypothesis. For example, isomerization of tau is positively correlated with levels of p62, a recognized indicator of autophagic inhibition. In sum, the data suggest strong ties between isomerization and autophagic flux, which may therefore represent a promising target for future investigations into the therapy and prevention of AD.

## Introduction

Despite decades of intense investigation, the underlying causes of Alzheimer’s disease (AD) remain incompletely understood. Many hypotheses have focused on aggregation of Aβ peptides, which form toxic oligomers as well as larger fibrils and ultimately extracellular deposits in multiple brain regions of people with AD.^1^ Abnormally phosphorylated and otherwise post-translationally modified tau has also received significant attention, as it is a primary constituent of neurofibrillary tangles (NFTs) and neuropil threads that are observed within neurons in partially overlapping brain regions of people with AD.^2^ The accumulation of pathologic tau is a better predictor of dementia,^3^ and recent work has shown that phosphorylated tau at positions 217, 181, and 205 indicate staging and development of AD^4^ with translation to reliable plasma markers that also correlate with degree of dementia.^5,6^ In addition, there are many other pathologic changes that accompany AD, including other aggregated deposits such as Lewy bodies,^7^ vascular disease,^8^ energy deficiency,^9^ and lysosomal storage.^10,11^ The relationships between all of these observations and their underlying causes remain to be fully elucidated, but it is clear that absent genetic predisposition, increasing age is the primary risk factor for AD.^12^

One aspect of aging that remains underexplored is the spontaneous chemical modification (SCM) of long-lived proteins. These modifications are not enzymatically catalyzed, and some fall largely outside the purview of biological control. Although many SCMs are easily detected and have been well documented, including oxidation,^13^ truncation,^14^ and deamidation,^15,16^ others are more pernicious and typically evade notice, such as isomerization.^17,18^ At the level of an individual residue, isomerization leads to dramatic structural changes, but these alterations are masked in the context of a protein where all other residues remain unmodified. Aspartic acid is the amino acid most prone to isomerize, producing four distinct structures by way of a succinimide intermediate.^19^ Although isomerization does not result in a change in mass, recently reported methods have enabled isomer detection in targeted proteomics-like experiments.^20,21^

In the more specific context of AD, isomerization has also received limited attention. Aβ extracted from amyloid plaques contains many sites of isomerization.^22,23,24^ This is perhaps not surprising, as extracellular plaques may persist for years, allowing ample opportunity for spontaneous modifications to occur. Isomerization of Aβ has been examined primarily within the context of its influence on aggregation, although recent results have illustrated that preventing lysosomal degradation may be more injurious.^25^ In the case of tau, isomerization has only been studied in a handful of reports. Initial work relied on methylation of L-isoAsp by protein isoaspartyl methyl-transferase (PIMT) to quantify isomerization of tau at the protein level.^26^ These results found greater isomerization in AD samples, but sites of modification and their relative contributions were not identified or quantified. Later work identified some sites of isomerization^27^ and examined qualitative spatial distributions by imaging select NFTs.^28^ However, despite the promising nature of these early results, many aspects of tau isomerization and its potential connection with AD remain unexplored.

In this study, we employ novel analysis of an emerging mass-spectrometry-based method to identify and quantify the extent of isomerization in different proteins extracted from human brain samples. We found striking differences in the amount of isomerization of tau between AD and control samples. Isomerization of tau also can differentiate sporadic versus autosomal dominant AD, with isomerization being greater in autosomal dominant. The data point towards defects in autophagic flux, which is strongly supported by additional correlations with other autophagic markers, including p62. The implications of these results in relation to potential treatment and therapy options for AD are discussed.

## Results

### DIA proteomics

To identify and quantify isomerization in human brain proteins, we carried out mass spectrometry-based proteomics experiments utilizing data-independent acquisition (DIA). In DIA, fragment-ion information is collected for all precursor ions present in a series of overlapping m/z ranges. Therefore, in theory, complete extracted ion chromatograms can be generated for all detectable peptide species. While isomers can theoretically be identified from these extracted ion chromatograms based on the presence of multiple peaks, typical DIA analysis pipelines only identify the best peak for each peptide.^29^ To overcome this limitation, the data was scanned for instances where two or more peaks eluted at different chromatographic retention times yet corresponded to the same peptide sequence based on high confidence matches of both precursor mass and numerous fragment ions. To eliminate false positives from partial matches of fragment ions that derived from nonequivalent precursors, the data from each peak was ranked against all potential precursors coeluting across both peaks. For non-isomeric peaks, this yields a higher score for different precursors for each peak whereas isomers yield similar scores for both peaks. We found 9 peptides from unique proteins meeting these criteria, which were assigned as potential sites of spontaneous isomerization, see Table S1.

Alternative explanations for the existence of such data were also considered but deemed unlikely. For example, for different retention times to result from switching the order of two residues by genetic mutation, the same mutation would have to be present in all of the different individuals who were sampled. Furthermore, although the precursor mass would be unchanged in such a circumstance, some fraction of the fragment ions would be able to distinguish such isomers. The peptides do not contain mass-shifting post-translational modifications (PTMs), so the results are not due to alternating PTM locations. Finally, exchange of Leu/IIe or vice versa is not possible in most cases as the peptides do not contain these residues. Therefore, the peptides eluting at multiple times most likely represent products of spontaneous isomerization. The degree of isomerization for each peptide can be calculated by comparing the relative abundances of the separate peaks, with the assumption that the most abundant peak (particularly in samples where one form is dominant) is the native form.

### Isomerization of Tau

Notably, tau is one of the isomer-containing proteins identified from the DIA analysis of identical amounts of total protein extracted from brain samples from 66 individuals. In Fig. 1a, the percent isomerization for the peptide ^386^TDHGAEIVYK^395^ is shown for 21 samples for sporadic AD and 2 samples for autosomal dominant AD (or ADAD) extracted from the hippocampus. The results are sorted by disease status and age, which is provided in the x-axis label. Although the percent isomerization values vary, it is clear that a considerable fraction of tau is isomerized in most AD/ADAD samples. In contrast, the % isomerization found in 22 hippocampal control samples was minimal (Fig.1b). The AD and controls are easily distinguished by statistical analysis (Fig. 1c), with averages of 14.4% and 1.9%, respectively. Note, the controls constitute two groups, control-high (12 samples) and control-low (10 samples), where control-high brains exhibited pathological features typically associated with AD but the person remained cognitively normal (additional detail is provided in Discussion and Supporting Information). Differences are more notable for samples extracted from the superior and middle temporal gyri (SMTG). In addition, the SMTG cohort contains 23 ADAD samples and 19 sporadic AD samples, allowing for statistical analysis of both cohorts. Most AD/ADAD samples exhibit some level of isomerization (Fig. 1d) while control samples (11 high, 9 low) are nearly devoid of isomerization (Fig. 1e). The magnitudes of the differences in the SMTG are also greater, yielding averages of 28.6%, 13.4%, and 0.33% isomerization for ADAD, sporadic AD, and control, respectively. The degree of isomerization in ADAD samples is distinguishable from both sporadic AD and controls (Fig. 1f). Notably, the majority of AD/ADAD samples exhibiting <5% isomerization in both the hippocampal and SMTG datasets are found in individuals over the age of 75 within the sporadic AD cohort. The degree of isomerization for the hippocampus and SMTG samples extracted from the same brains show strong correlation (Pearson’s 0.85), after exclusion of one outlier (Fig. S1). In contrast, other proteins identified in Table S1 do not exhibit different degrees of isomerization between AD and control groups.

**Figure 1.**
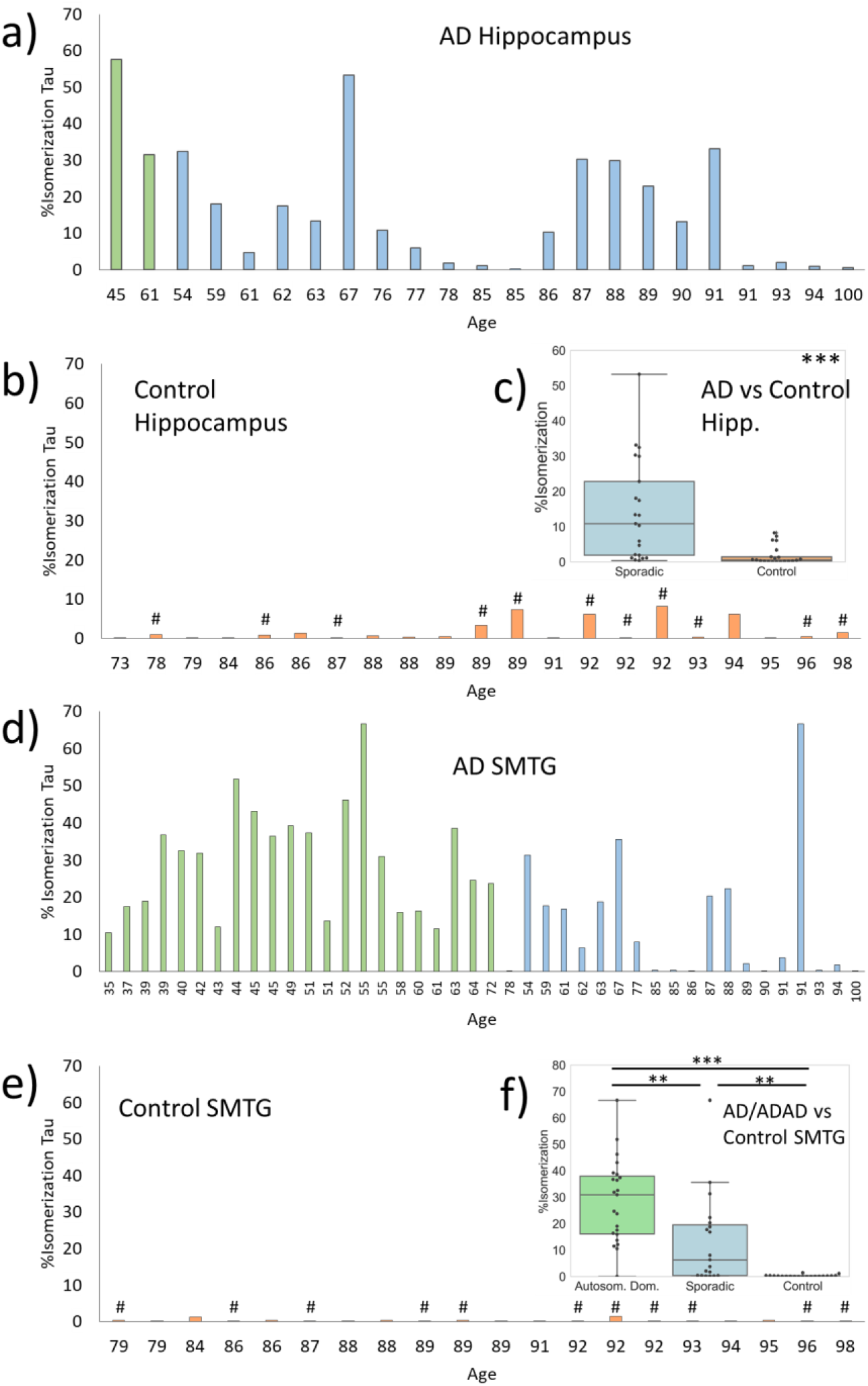
%isomerization of Asp387 in tau for a) AD (blue) and ADAD (green) samples, and b) control samples (orange) extracted from hippocampus. c) Comparison of %isomerization in AD versus control (note: ADAD values are not included). d)-f) analogous plots for SMTG. # indicates control-high. P-values are indicated by: * <0.05, ** <0.01, ***, <0.001.

To confirm the identity of isomers in the tissue samples, we collected LC-MS/MS data for a set of known synthetic tau isomer standards. Comparison of relative elution times confirms that ^386^TDHGAEIVYK^395^ from tau is isomerized at Asp387, in agreement with known amino acid propensities for isomerization and previous examination of tau, see Fig. S3. Synthetic standards also confirm the initial assignment of the native L-Asp form as the most abundant peak, followed by L-isoAsp. This observation is in agreement with previous results showing that L-isoAsp is the primary product of Asp isomerization.^30^

### Autophagy and Repair

In addition to our analysis of isomerization, we also quantified the amount of 13 proteins connected to autophagy using traditional DIA methodology. Seven (MPCP, MLP3A(LC3), Hsc70, Mtp70, p62, progranulin, and mTOR) exhibited some differences between AD and control groups, and six could not be distinguished between AD and control groups (Lamp2, CatB, Rack1, Hsp70, BiP, and 28S-RP-S27). Most strikingly, the chaperone p62 (a known marker for autophagy),^31^ shares the closest relationship with isomerization of tau, see Fig. 2. The relative abundance of p62 (represented by a z-score) correlates with the degree of isomerization of tau in both the hippocampus and SMTG AD/ADAD samples, yielding Pearson correlations of 0.89 for the hippocampus and 0.76 for the SMTG. In contrast, p62 levels do not correlate with isomerization in controls. The total amount of p62 is also higher in AD/ADAD samples, particularly in the SMTG (Fig. 2c). The levels of several other proteins connected with autophagy (MPCP, MLP3A, and Hsc70) were also evaluated and found to be reduced by modest amounts in AD (Fig. 2c).

**Figure 2.**
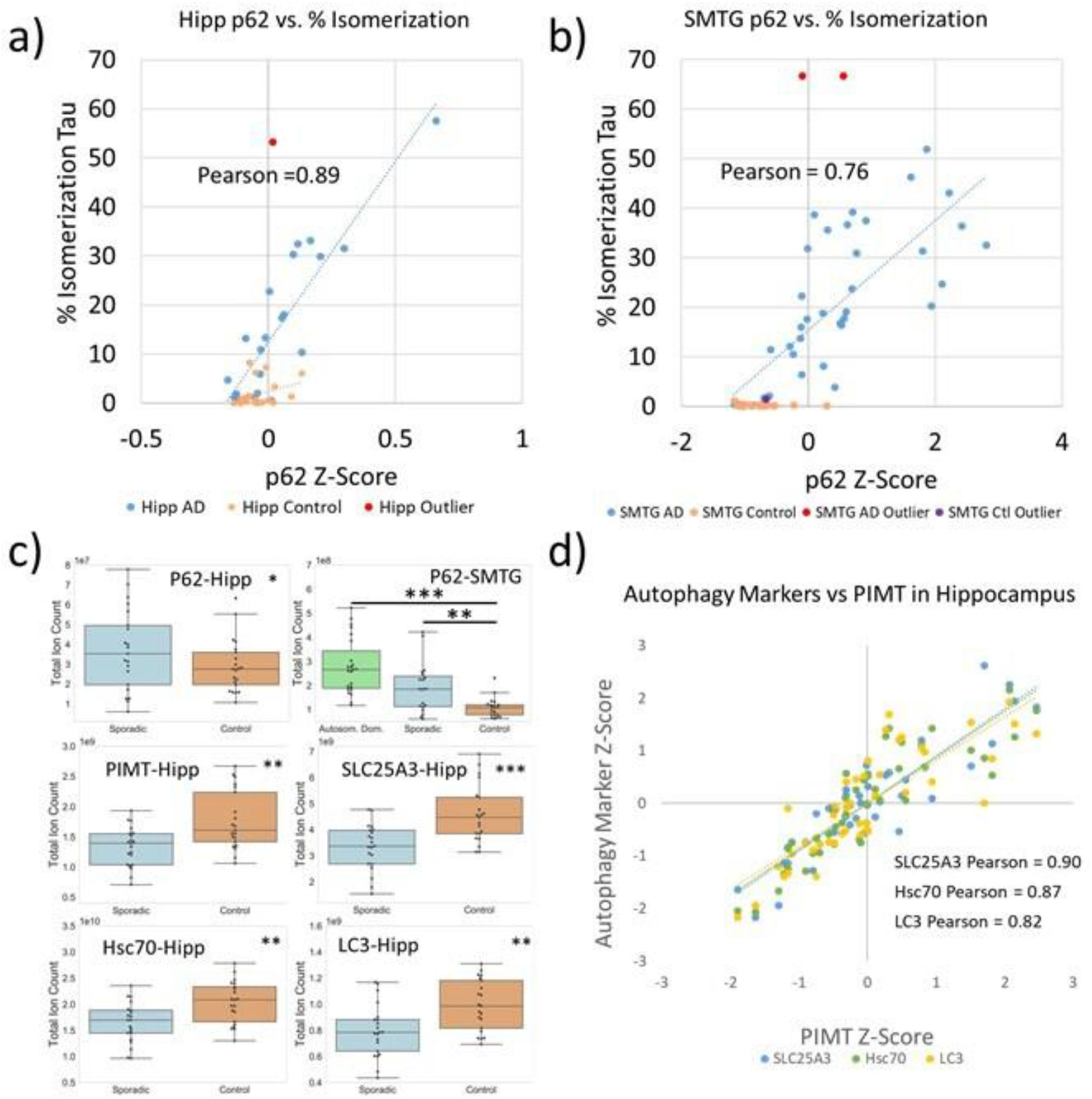
Correlation of tau %isomerization with p62 levels in the a) hippocampus and b) SMTG. c) Statistical comparisons of protein levels present in AD and control samples. Hippocampal samples do not include ADAD results. d) Correlation between various autophagy markers and PIMT levels.

Protein isoaspartyl methyl transferase (PIMT) is a repair enzyme that partially restores L-isoAsp to L-Asp. PIMT was detected in our dataset, and quantitative analysis revealed lower amounts of PIMT in hippocampal AD samples (Fig. 2c). Correlation between PIMT levels and isomerization of tau in the hippocampal dataset was modest (Pearson’s −0.39). Interestingly, the levels of MPCP, MLP3A, and Hsc70 correlate more strongly with PIMT levels (Fig. 2d), with Pearson values above 0.80.

## Discussion

The results in Fig. 1 clearly illustrate that the % isomerization of tau is a strong indicator of AD/ADAD status. In the hippocampus samples, tau isomerization is 8x greater in AD relative to controls. In the SMTG dataset, tau isomerization is 87x greater in ADAD and 41x greater in sporadic AD relative to controls. One important difference between the hippocampus and SMTG results is the near absence of isomerization in SMTG control samples. Although the overall trend is the same in both the hippocampus and SMTG, it is clear that different regions of the brain do not behave identically with regard to tau isomerization. Importantly, there are two separate control groups in our analysis (denoted as control-high and control-low). The control-low group consists of individuals with limited observable pathology (B scores of 2 or lower and C scores of 0) who scored >90 on the CASI (Cognitive Abilities Screening Instrument) test within two years of death. In contrast, the control-high group is unusual and constitutes individuals with B scores of 2/3 and C scores of 2/3, yet also scored >90 on CASI within two years of death. In other words, in terms of B and C scores, the control-high group is very similar to the AD and ADAD cohorts, yet they did not exhibit signs of dementia within two years of death. Strikingly, isomerization of tau is not abundant in either control group (or distinguishable between control-low and control-high), meaning that isomerization can clearly distinguish control-high from AD/ADAD where B/C scores cannot. This suggests that tau isomerization occurs independently from aberrant protein aggregation, yet it is related in some fashion to the processes leading to cognitive and functional impairment.

Isomerization of amino acid residues in proteins is a spontaneous chemical modification that requires no enzymatic intervention to proceed.^32^ The isomerization we observe in tau occurs primarily at Asp387, consistent with previous studies showing that Asp is the most readily isomerized amino acid in long-lived proteins^,19,20,22^ and that DH motifs are prone to isomerize.^30^ Because the process is spontaneous, for significant amounts of isomerization to accumulate in the brain, protein turnover must be sufficiently slow to allow isomerization to occur. Although the isomerization rate of Asp387 in tau is not known, measured rates of Asp isomerization in other peptides were found to proceed at rates of ~1% per day or less.^25^ If isomerization of Asp387 in tau proceeds similarly, then the amount of isomerization reported in Fig. 1 would require up to 60 days (or more) for the most dramatic cases. Previously measured lifetimes for soluble tau exceeded 60 days for the fraction secreted into cerebrospinal fluid,^33^ which is consistent with this timeframe. Importantly, soluble tau is intrinsically disordered,^34^ which is known to facilitate isomerization of Asp residues in general. This property of tau may facilitate isomerization on a timescale relevant to AD pathology.

We hypothesize that the increased tau isomerization in AD reflects a slowdown of autophagy, which increases tau lifetime and decreases its turnover rate (allowing for greater isomerization to occur). To support this hypothesis, we examined the levels of autophagic markers within our DIA dataset. A variety of autophagy related proteins are present in lower amounts in the AD cohorts, see Fig. 2c.^35,36,37^ Although the differences are statistically significant, the magnitudes of the changes are not particularly large. However, we did discover greater differences in one autophagic marker, p62, which is known to accumulate when autophagy is impaired.^31^ The amount of p62 is greater in AD by a factor of 1.6x in the hippocampus and 2.2x in the SMTG. Although previous studies of the frontal cortex using Western blot found lower levels of p62 in AD brains relative to controls, those results may represent levels of cytosolic p62.^38^ Consistent with this possibility, other studies have demonstrated strong co-localization of p62 within tau inclusions.^39^ In our studies, we found strong correlation between the amount of p62 and degree of isomerization in samples for both the hippocampal and SMTG datasets (Fig. 2a and 2b), although isomerization more clearly distinguished AD/ADAD samples from controls. The buildup of p62 supports the idea that isomerization is greater in AD/ADAD due to slowed autophagic flux, which is known to cause p62 accumulation,^40^ although there are known limitations to the use of p62 to monitor autophagic flux.^41^ Overall, our findings also support other connections that have been reported between autophagy, p62, and AD.^42,43^

It has been suggested previously that tau isomerization may simply be a consequence of the long half-life of tau present in neurofibrillary tangles (NFTs),^27,28^ but our data is not consistent with this interpretation. For example, isomerization is nearly absent in the control-high group and some AD samples despite the presence of abundant NFTs in both. Furthermore, comparison of tau isomerization vs age within the hippocampal sporadic AD cohort revealed poor correlation (Pearson −0.35) and the trend is downward with age. To be consistent with isomerization representing NFT half-life, this would require NFTs in older people to have been present for less time. Furthermore, continued isomerization may not be possible for tau trapped in NFTs, in which case NFT age would not correlate with degree of isomerization. Indeed, structures for the fibrils constituting neurofibrillary tangles have recently been determined, and Asp387 lies just outside the highly ordered domain in a partially ordered region.^44^ It may be difficult to accommodate succinimide formation in such close proximity to densely packed fibrils where the structural flexibility required for isomerization to occur is likely to be hindered. Finally, recent results have demonstrated that tau exchanges dynamically in and out of NFTs,^45^ which further complicates the relationship between tau and NFT lifetimes. Similar arguments apply to the possibility that degree of isomerization might represent overall NFT abundance. Since isomerization decreases with age, the abundance of NFTs would need to decrease on average for older individuals, which has not been the observed trend.^46^ Given the totality of these considerations, it is likely that our results primarily reflect the extent of isomerization in ‘distributed’ tau, or tau that is *not present* in NFTs. This could include fully functional tau, smaller soluble aggregates, or undigested tau trapped in autophagic vacuoles or other lysosomal bodies.

To ascertain the extent to which other important factors might influence the amount of tau isomerization, we examined the amount of the repair enzyme PIMT in each sample. PIMT can repair some products of Asp isomerization by methylation, which occurs primarily at L-isoAsp.^47,48^ Comparison between AD and controls reveals that the levels of PIMT are lower in AD samples, although the effect is modest and unlikely to fully account for the observed differences in isomerization. In addition, there is weak correlation between PIMT levels and isomerization of tau, further suggesting that PIMT is not a dominant controlling factor. However, levels of some of the autophagic markers in Fig. 2 do show strong correlation with the level of PIMT (Fig. 2d), suggesting that expression of PIMT is connected with (or perhaps regulated by) autophagy.

### Connections to autophagy

The disease-causing mutations within our ADAD cohort occur exclusively at *PSEN1* and *PSEN2*. The associated proteins, presenilin 1 (PS1) and presenilin 2 (PS2), function as components of γ-secretase complex, which is responsible for enzymatic cleavage of amyloid precursor protein.^49^ Although mutant presenilins are often discussed in the context of Aβ cleavage,^50^ the same modifications can also impact autophagy.^51,52^ For example, mutant presenilins are associated with increased lysosomal malfunction that may occur by disruption of lysosomal acidification or autophagosome formation.^53,54^ In other cases, evidence implicates disruption of calcium homeostasis as the source of lysosomal impairment.^55,56^ Regardless of the precise underlying cause, there is a clear potential for disrupted autophagy resulting from the mutations in *PSEN1* and *PSEN2*. Our data reveals that tau isomerization is greatest in ADAD (Fig. 1f), suggesting that autophagic flux is reduced most in ADAD. The lesser (yet still greater than control) degree of isomerization in sporadic AD suggests a more subtle diminution of autophagic flux. Consistent with this thought, numerous studies have shown that autophagy declines with age in general.^57,58^ Furthermore, PS1 expression naturally declines with increasing age.^52^ Decreased autophagic flux is therefore consistent with age being the greatest risk factor in AD.

## Conclusion

Collectively, our observations point towards a connection between autophagic flux and isomerization of tau. Increased isomerization observed in a broad collection of samples suggests that autophagy is impaired in both the autosomal dominant and sporadic forms of the disease. Necessarily, lower autophagic flux would lead to reduction in the clearance of proteins such as tau and Aβ. Much has been discussed about the aggregation propensities of these species, for example that Aβ 1-42 aggregates much more readily than Aβ 1-40.^1^ Or that seed forms of tau can propagate aggregation.^59,60^ However, tau and Aβ are both present from birth, yet apparently remain unproblematic for decades. A trigger that corresponds with aging is required to ‘activate’ the pathogenic properties of Aβ and tau. Our isomerization data suggests that reduced autophagic flux may be this trigger. The additional protein lifetime that follows this slowdown affords the opportunity not only for isomerization, but also other deleterious modifications, including aggregation or hyper-phosphorylation. In other words, frequent turnover may be the primary cellular mechanism for preventing isomerization or aggregation from becoming pathological. Furthermore, once autophagic flux slows sufficiently to allow isomerization, a destructive cycle is enabled because isomerization itself is refractory to and interferes with autophagic degradation.^25^

## Acknowledgements

The authors gratefully acknowledge funding from the NIH (R01 AG066626 for RRJ, RF1 AG053959 for MM and TJM, U01 AG006781 (ACT study) and P30 AG066509 (UW ADRC), and the Nancy and Buster Alvord Endowment (CDK), P30 AG062421 for MPF, U19 AG032438 (DIAN) for RB. Data collection and sharing for this project was supported by The Dominantly Inherited Alzheimer’s Network (DIAN, UF1AG032438, see ListS1 in supporting information) funded by the National Institute on Aging (NIA), the German Center for Neurodegenerative Diseases (DZNE), Raul Carrea Institute for Neurological Research (FLENI), Partial support by the Research and Development Grants for Dementia from Japan Agency for Medical Research and Development, AMED, and the Korea Health Technology R&D Project through the Korea Health Industry Development Institute (KHIDI). This manuscript has been reviewed by DIAN Study investigators for scientific content and consistency of data interpretation with previous DIAN Study publications. We acknowledge the altruism of the participants and their families and contributions of the DIAN research and support staff at each of the participating sites for their contributions to this study.

## Author Contributions

Conceptualization, RRJ and MM; Methodology, RRJ, MM, EEH, LH, GEM; Investigation, EEH, LH, GEM; Tissue processing, JPC, MRF,CAM, BG, KLN, MPF, RJB, EBL, CDK; Writing-original draft, RRJ, EEH, LH; Writing-reviewing/editing; all authors.

## Methods

### Sample Stratification Description

Brain tissue samples were stratified into 4 groups based on clinical, pathological and genetic data (Table S2A and S2B). Cognitive status was determined as dementia (AD/ADAD) or no dementia (controls) by DSM-IVR criteria; controls were from the Adult Changes in Thought (ACT) study and were included only if the last evaluation was within 2 years of death and the last CASI score was >90 (upper quartile for controls in the ACT cohort). Controls with no or low Alzheimer’s disease neuropathologic change (ADNC) were designated ‘control-low’ and those with intermediate or high ADNC were designated ‘control-high’. All cases with dementia had intermediate or high level ADNC and were classified as AD, and further subclassified as sporadic AD or autosomal Dominant AD (ADAD) when a dominantly inherited mutation in *PSEN1* or *PSEN2* was found. Sporadic AD cases came from the ACT and the University of Washington (UW) AD Research Center (ADRC), whereas ADAD cases came from the UW ADRC and the Dominantly Inherited Alzheimer Network (DIAN). Cases with Lewy body disease (other than those involving only the amygdala), territorial infarcts, more than 2 cerebral microinfarcts, or hippocampal sclerosis were excluded. Time from death to cryopreservation of tissue, post mortem interval (PMI), was <8 hr in all cases except for those in the ADAD cohort. Details of sample stratification for the 2 brain regions (SMTG and Hippocampus) are provided in Tables S2A and S2B in the supporting information.

### Sample Metadata, Batch Design and References

Each region of human brain tissue was divided into batches of 14 individual samples and 2 pooled references for a total of 16. The first batch of each region was also used to create a region-specific reference pool to be used as a “common reference” and/or single point calibrant, which was homogenized, aliquoted, frozen, and used to compare between batches within a brain region. Human cerebellum and occipital lobe tissue was homogenized, pooled, aliquoted and frozen to be used as a “batch reference” for comparison between batches and other brain regions. Batch design was randomly balanced based on group ratios. For example, batches from the SMTG brain region contained 5 “Sporadic AD”, 4 “Autosomal Dominant AD”, 2 “Control Low Neuropathology”, and 3 “Control High Neuropathology” samples.

### Lysis/Digestion

Two 25 μm cryo slices (“curls”) of brain tissue were resuspended in 120 μl of lysis buffer of 5% SDS, 50mM Triethylammonium bicarbonate (TEAB), 2mM MgCl2, 1X HALT phosphatase and protease inhibitors, vortexed, and briefly sonicated at setting 3 for 10 seconds with a Fisher sonic dismembrator model 100. A microtube was loaded with 30 μl of lysate and capped with a micropestle for homogenization with a Barocycler 2320EXT (Pressure Biosciences Inc.) for a total of 20 minutes at 35°C with 30 cycles of 20 seconds at 45,000 psi followed by 10 seconds at atmospheric pressure. Protein concentration was measured with a BCA assay. Homogenate of 50 μg was added to a process control of 800 ng of yeast enolase protein (Sigma) which was then reduced with 20 mM DTT and alkylated with 40 mM IAA. Lysates were then prepared for S-trap column (Protifi) binding by the addition of 1.2% phosphoric acid and 350 μl of binding buffer (90% Methanol, 100 mM TEAB). The acidified lysate was bound to column incrementally, followed by 3 wash steps with binding buffer to remove SDS and 3 wash steps with 50:50 methanol:chloroform to remove lipids and a final wash step with binding buffer. Trypsin (1:10) in 50mM TEAB was then added to the S-trap column for digestion at 47°C for one hour. Hydrophilic peptides were then eluted with 50 mM TEAB and hydrophobic peptides were eluted with a solution of 50% acetonitrile in 0.2% formic acid. Elutions were pooled, speed vacuumed and resuspended in 0.1% formic acid.

Injection of samples consisted of one μg of total protein (16 ng of enolase process control) and 150 fmol of a heavy labeled Peptide Retention Time Calibrant (PRTC) mixture (Pierce). The PRTC was used as a peptide process control. Library pools contained an equivalent amount of every sample (including references) in the batch. For example, a batch library pool consisted of the 14 samples from the batch and two references. System suitability (QC) injections comprised 150 fmol of PRTC and BSA.

### Liquid Chromatography and Mass Spectrometry

One μg of each sample with 150 femtomole of PRTC was loaded onto a 30 cm fused silica picofrit (New Objective) 75 μm column and 3.5 cm 150 μm fused silica Kasil1 (PQ Corporation) frit trap loaded with 3 μm Reprosil-Pur C18 (Dr. Maisch) reverse-phase resin analyzed with a Thermo Easy-nLC 1200. The PRTC mixture was used to assess system suitability before and during analysis. Four of these system suitability runs were analyzed prior to any sample analysis and then after every six sample runs another system suitability run was analyzed. The 110-minute sample LC gradient consists of a 2 to 7% for 1 minutes, 7 to 14% B in 35 minutes, 14 to 40% B in 55 minutes, 40 to 60% B in 5 minutes, 60 to 98% B in 5 minutes, followed by a 9 minute wash and a 30 minute column equilibration. Peptides were eluted from the column with a 50°C heated source (CorSolutions) and electrosprayed into a Thermo Orbitrap Fusion Lumos Mass Spectrometer with the application of a distal 3 kV spray voltage. For the sample digest, first a chromatogram library of 6 independent injections was analyzed from a pool of all samples within a batch. For each injection a cycle, one 120,000 resolution full-scan mass spectrum was acquired with a mass range of 100 *m/z* (400-500 *m/z*, 500-600 *m/z*…900-1000 *m/z*) followed by data-independent MS/MS spectra on the loop count of 26 at 30,000 resolution, AGC target of 4e5, 60 sec maximum injection time, 33% normalized collision energy with a 4 *m/z* overlapping isolation window.^61,62,63^ The chromatogram library data was used to quantify proteins from individual sample runs. These individual runs consisted of a cycle of one 120,000 resolution full-scan mass spectrum with a mass range of 350-2000 *m/z*, AGC target of 4e5, 100 ms maximum injection time followed by a data-independent MS/MS spectra on the loop count of 76 at 15,000 resolution, AGC target of 4e5, 20 sec maximum injection time, 33% normalized collision energy with an overlapping 8 *m/z* isolation window. Application of the mass spectrometer and LC solvent gradients were controlled by the ThermoFisher XCalibur (version 3.1.2412.24) data system.

### Data Analysis

Thermo RAW files were converted to mzML format using Proteowizard (version 3.0.20064) using vendor peak picking and demultiplexing.^64^ Chromatogram spectral libraries were created using default settings (10 ppm tolerances, trypsin digestion, HCD b- and y-ions) of of EncyclopeDIA (version 0.9.5) using a Prosit predicted spectra library based the Uniprot human canonical FASTA.^65^ Quantitative spectral libraries were created by mapping spectra to the chromatogram spectral library using EncyclopeDIA^66,67,68^ requiring a minimum of 3 quantitative ions and filtering peptides at a 1% FDR using Percolator 3.01.^69,70^ The quantitative spectral library is imported into Skyline (daily version 20.1.1.83) with the human uniprot FASTA as the background proteome to map peptides to proteins.^71,72^ Transition retention time filtering settings set to “use only scans within 4 minutes of MS/MS IDs” for all batches and then removed all peptides/proteins from Skyline document except APP (Uniprot Accession P05067) and Tau (Uniprot Accession P10636).

### Automated identification of potential isomers

Raw files collected with 4-m/z staggered isolation windows were demultiplexed and converted to mzMLs using MSConvert (Proteowizard, v 3.0.20239).^73^ The resulting mzMLs were searched in Crux Tide (v 3.2) and the top 10,000 precursors per spectrum were reported so that the results file contained every possible spectrum/ peptide combination.^74^ To limit search size, two searches were performed. The first search covered the entire tryptic human proteome (UP000005640) with no allowed missed cleavages or modifications. The second search covered a small subset of relevant proteins allowing semi-tryptic peptides with up to two missed cleavages, N-terminal acetylation, and methionine oxidation. All peptides and spectra from each 2-m/z isolation window were converted into matrices with Tailor calibrated XCorr as the values where each row is a potential peptide and each column is a spectrum.^75,76^ All candidate peptides were filtered so they had at least one score above a given noise threshold, here we used a cutoff of 1.15 for the full proteome search and 1.1 for the smaller semi-tryptic search. Scores were baseline adjusted with simple subtraction so that the minimum score for each peptide was 0. Peaks were defined as multiple consecutive spectra scoring above 80% of the maximum baseline-subtracted score. The remaining candidates were filtered for peptides with multiple peaks separated by 4 or more scans below the 80% threshold. With the remaining list of peaks, overlapping peaks that were generated by two different peptides were identified. In each of these cases, angle cosine was calculated between the lower scoring of the two peptides and 1000 random decoys as well as the lower scoring and higher scoring peptide. If the angle cosine between the two candidate peptides was greater than the maximum angle cosine between the decoys, it is likely that these peaks were caused by the same sets of ions. Therefore, the lower scoring of the two peaks was removed and the process was repeated for all identified peaks. All peptides with two or more peaks remaining after this process were identified as likely stereoisomers.

### Quantification of isomerization and protein biomarkers

Isomer quantification was determined using peak area values extracted from Skyline (Skyline-daily 20.9.234). To calculate peak areas, the boundaries for each individual isomer peak were adjusted manually to account for differences in retention times between chromatograms. Within this window, the total area of both the precursor and fragment peaks corresponding the peptide of interest were then summed to produce a “total isomer area” value for each individual isomer. Fragments or precursors not found in all datasets were not used to calculate area values to prevent skewing the results with marginally detectable ions. Using total isomer area values, the “percent isomerization” was calculated by dividing the sum of isomer peak areas by the total peak area (where the canonical form was assumed to be dominant), see eq. 1. The ionization efficiencies of isomeric peptides are expected to be very similar due to identical size, functional group composition, and similar structure. Furthermore, any differences in ionization would be reproduced between sample sets, meaning that comparisons of differences between samples will accurately reflect differences in the amounts of isomers present. Means, two-tailed p-values, and correlations were calculated in Microsoft Excel.

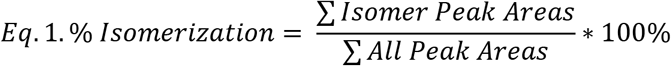

Protein quantification was determined for each sample by using the sum of all detected peptide areas assigned within the chromatogram by Skyline (eq. 2). The reported values therefore represent relative abundance normalized to the total ion count. Peptides not found in all datasets were not used to calculate protein area values. It was assumed that differences in fragment ions between datasets would not be significant, due to dilution across the large number of discovered peptides for each protein. Means, two-tailed p-values, correlations, and z-score conversions were calculated in Excel.

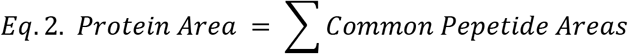

### Peptide synthesis and isomer verification

Peptide standards for each isomeric form were synthesized following a standard solid phase peptide synthesis protocol.^77^ The synthesized standards were then lyophilized to dryness, dissolved in 1mL 50:50 ACN:H_2O_, and frozen in a −20C freezer. Sequence was confirmed by electrospraying a 5uM peptide in H_2O_ with 0.1% formic acid to verify exact mass and fragmentation patterns consistent with the desired peptide. An isomer mixture was then run through a setup equivalent to that used for all DIA data collection.

### Data Availability

The Skyline documents and raw files for mass spectrometry data will be available at Panorama Public.^78,79^

