## supporting information for "Spontaneous Isomerization of Asp387 in Tau is Diagnostic for Alzheimer’s Disease: An Endogenous Indicator of Reduced Autophagic Flux"

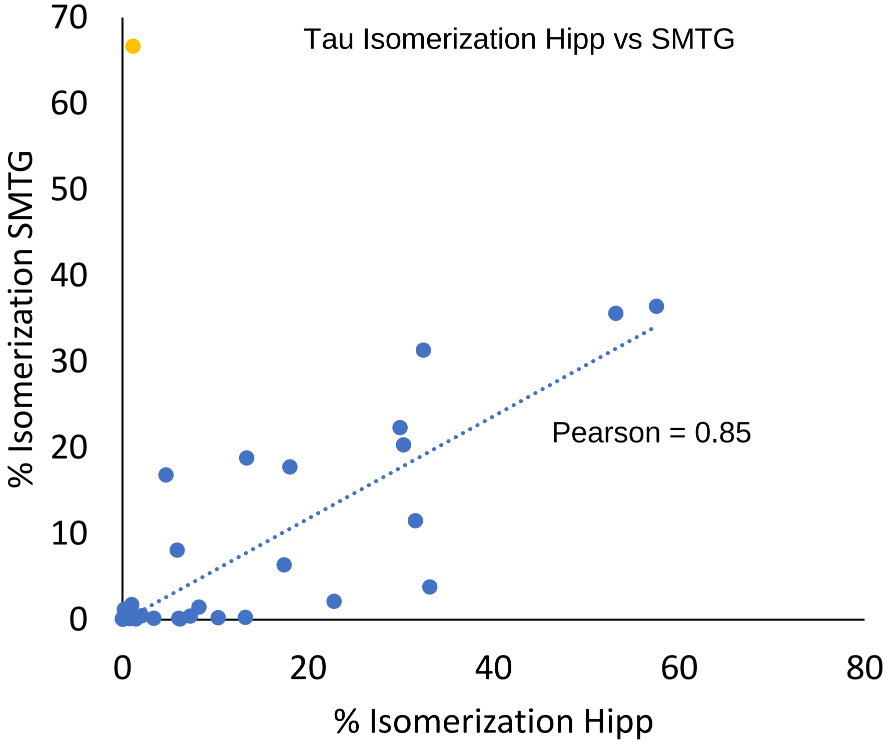


**Figure S1.** Comparison of Tau isomerization in the Hippocampus and SMTG regions in brains from which both regions were examined. A single outlier exhibits high levels of SMTG isomerization concurrent with virtually no isomerization in the hippocampus.

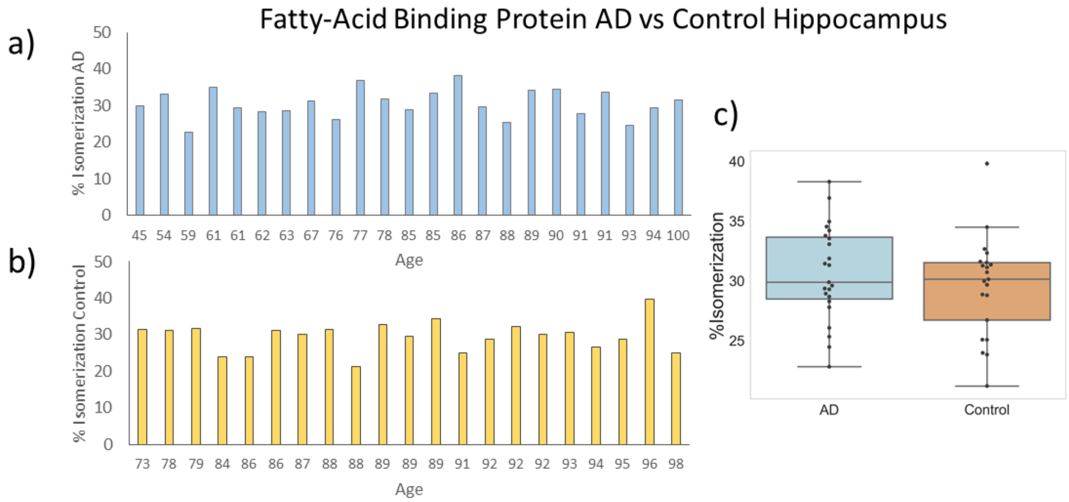


**Figure S2.** % Isomerization of the discovered isomerized peptide WDGQETTLVR [97, 106] of *fatty-acid binding protein, heart* in hippocampus. (a) % Isomerization of the AD group, samples organized by age. (b) % Isomerization of the control group. (c) Boxplot of % isomerization of AD and Control groups. P=0.40. Extent of isomerization is consistent across AD and control populations.

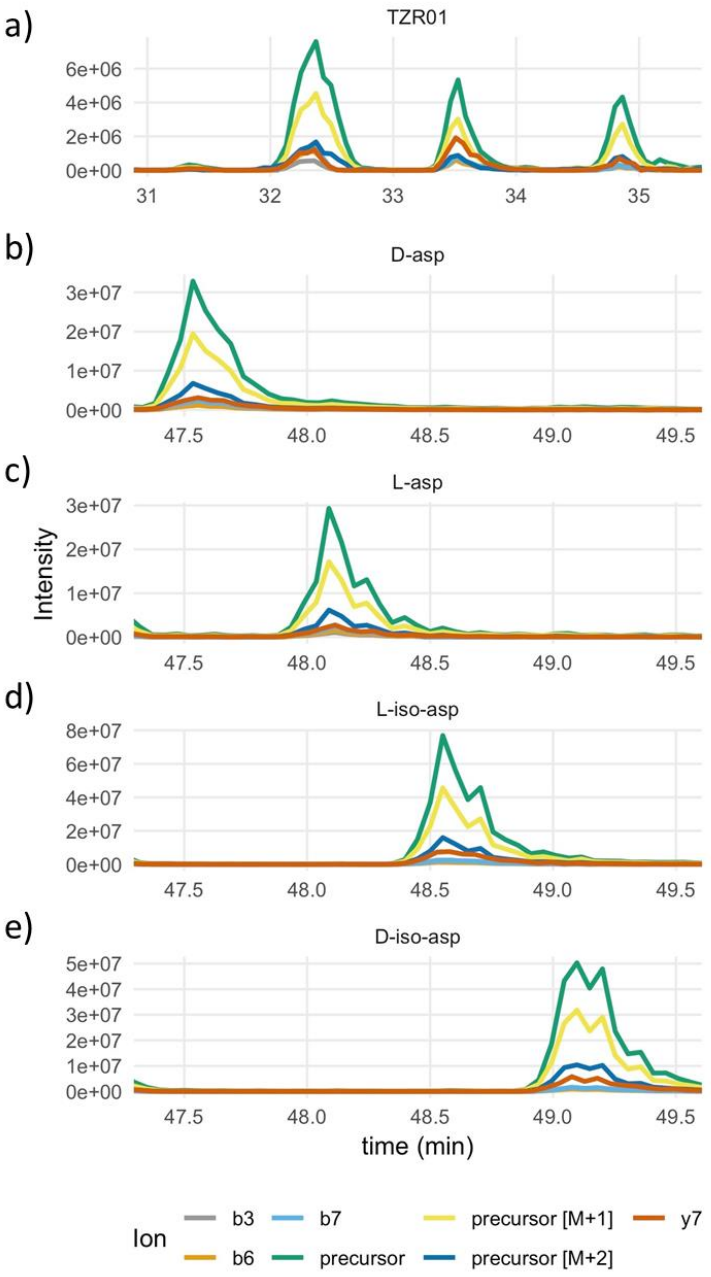


**Figure S3.** Comparison of DIA chromatogram of TDHGAEIVYK from the SMTG region (a) to chromatograms of synthetic standards of TDHGAEIVYK aspartic acid (asp) isomers (b-e). Elution orders indicate that peaks observed in DIA data are, in order of retention time, D-asp, L-Asp, L-isoAsp, and D-isoAsp.

**Table S1. Discovered Isomerized Proteins in Hippocampus**

| Protein | Peptide | AD Mean | Control Mean | P-Value |
|---|---|---|---|---|
| Microtubule-associated protein tau | TDHGAEIVYK [702,711] | 17.1% | 1.86% | 0.00027 |
| Sodium/calcium exchanger 2 | AAPAEGAGEDEDDGASR [378, 394] | 11.8% | 11.0% | 0.51 |
| Protein NipSnap homolog 3A | QYDGIFYEFR [30, 39] | 25.5% | 25.8% | 0.90 |
| Tubulin beta-3 chain | MSSTFIGNSTAIQELFK [362, 378] | 38.4% | 36.9% | 0.73 |
| Spectrin beta chain, non-erythrocytic 1 | FATDGEGYKPCDPQVIR [593, 609] | 16.1% | 19.7% | 0.06 |
| Receptor-type tyrosine-protein phosphatase zeta | FAVLYQQLDGEDQTK [345, 359] | 40.9% | 39.5% | 0.49 |
| Annexin A2 | AEDGSVIDYELIDQDAR [179, 195] | 28.9% | 32.4% | 0.22 |
| Fatty acid-binding protein, heart | WDGQETTLVR [97, 106] | 32.3% | 31.0% | 0.40 |
| N(G),N(G)-dimethylarginine dimethylaminohydrolase 1 | DYAVSTVPVADGLHLK [159, 174] | Isomer peak is cut off in Skyline, not analyzed | | |

**Table S1.** Proteins discovered to contain isomerized peptides. Excluding tau, means and p-values were determined using a limited dataset of values (AD n=10, Control n=6) from hippocampus samples. No proteins aside from tau were statistically different in AD vs. Control. *Spectrin beta chain, non-erythrocytic 1* appeared to have potential statistical differences, but examining the entire hippocampus dataset revealed p=0.51. In *N(G),N(G)-dimethylarginine dimethylaminohydrolase 1*, an isomer peak fell partially outside of the chromatogram window, and a % isomerization value could not be accurately calculated.

**Table S2A. Brain Tissue Stratification Description for SMTG**

| | Control Low Path | Control High Path | Sporadic AD | Autosomal Dominant AD |
|---|---|---|---|---|
| N | 9 | 11 | 19 | 23 |
| Age (yr)* | 88 + 5 | 90 + 5 | 80 + 14 | 51 + 11 |
| Sex (M:F) | 4:5 | 5:6 | 11:8 | 15:8 |
| Post mortem interval (hr)* | 3.9 + 0.9 | 4.9 + 1.5 | 4.6 + 1.2 | 16.2 + 9.3 |
| # *APOE* ε4 alleles | 2 of 18 | 6 of 22 | 9 of 38 | 4 of 32 |
| *PSEN1* mutations | None | None | None | Y115C, 2x A260V, G206V, I229F, M233L, 3 x G209V, 2 xI143T, N135S, T245P, 2 x H163R, A431E, S169L |
| *PSEN2* mutations | None | None | None | 6 x N141I |
| B Score (0 to 3 scale) | 4 x B1, 5 x B2 | 5 x B2, 6 x B3 | B3 in all cases | B3 in all cases |
| C Score (0 to 3 scale) | C0 in all cases | 5 x C2, 6 x C3 | 3 x C2, 16 x C3 | 1 x C2, 22 x C3 |

*mean $\pm$ SD

**Table S2B. Brain Tissue Stratification Description for Hippocampus**

| | Control Low Path | Control High Path | Sporadic AD | Autosomal Dominant AD |
|---|---|---|---|---|
| N | 10 | 11 | 21 | 2 |
| Age (yr)* | 87 + 6 | 90 + 5 | 80 + 13 | 53 + 8 |
| Sex (M:F) | 5:5 | 5:6 | 12:9 | 2:0 |
| Post mortem interval (hr)* | 3.9 + 0.9 | 5.1 + 1.3 | 4.5 + 1.3 | 9.8 + 6.2 |
| # *APOE* ε4 alleles | 2 of 20 | 6 of 22 | 12 of 42 | 0 of 4 |
| *PSEN1* mutations | None | None | None | G209V, A431E |
| *PSEN2* mutations | None | None | None | None |
| B Score (0 to 3 scale) | 5 x B1, 5 x B2 | 5 x B2, 6 x B3 | B3 in all cases | B3 in all cases |
| C Score (0 to 3 scale) | C0 in all cases | 5 x C2, 6 x C3 | 4 x C2, 17 x C3 | C3 in all cases |

*mean ± SD

**ListS1: Full Name and Credentials of the Dominantly Inherited Alzheimer Network**

Sarah Adams, MS; Ricardo Allegri, PhD; Aki Araki, ; Nicolas Barthelemy, PhD; Randall Bateman,MD; Jacob Bechara, BS; Tammie Benzinger, MD, PhD; Sarah Berman, MD, PhD; Courtney Bodge, PhD; Susan Brandon, BS; William (Bill) Brooks, MBBS,MPH; Jared Brosch, MD, PhD; Jill Buck, BSN; Virginia Buckles, PhD; Kathleen Carter, PhD; Lisa Cash, BFA; Charlie Chen, BA; Jasmeer Chhatwal, MD,PhD; Patricio Chrem Mendez, MD; Jasmin Chua, BS; Helena Chui, MD; Laura Courtney, BS; Carlos Cruchaga, PhD; Gregory S Day, MD; Chrismary DeLaCruz, BA; Darcy Denner, PhD; Anna Diffenbacher, MS; Aylin Dincer, BS; Tamara Donahue, MS; Jane Douglas, MPh; Duc Duong, BS; Noelia Egido, BS; Bianca Esposito, BS; Anne Fagan, PhD; Marty Farlow, MD; Becca Feldman, BS,BA; Colleen Fitzpatrick, MS; Shaney Flores, BS; Nick Fox, MD; Erin Franklin, MS; Nelly Joseph-Mathurin, PhD; Hisako Fujii, PhD; Samantha Gardener, PhD; Bernardino Ghetti, MD; Alison Goate, PhD; Sarah Goldberg, MS,LPC,NCC; Jill Goldman, MS,MPhil,CGC; Alyssa Gonzalez, BS; Brian Gordon, PhD; Susanne Gräber-Sultan, PhD; Neill Graff-Radford, MD; Morgan Graham, BA; Julia Gray, MS; Emily Gremminger, BA; Miguel Grilo, MD; Alex Groves, ; Christian Haass, PhD; Lisa Häsler, MSc; Jason Hassenstab, PhD; Cortaiga Hellm, BA; Elizabeth Herries, BA; Laura Hoechst-Swisher, MS; Anna Hofmann, MD; Anna Hofmann, ; David Holtzman, MD; Russ Hornbeck, MSCS, MPM; Yakushev Igor, MD; Ryoko Ihara, MD; Takeshi Ikeuchi, MD; Snezana Ikonomovic, MD; Kenji Ishii, MD; Clifford Jack, MD; Gina Jerome, MS; Erik Johnson, MD, PHD; Mathias Jucker, PhD; Celeste Karch, PhD; Stephan Käser, PHD; Kensaku Kasuga, MD; Sarah Keefe, BS; William (Klunk, MD, PHD; Robert Koeppe, PHD; Deb Koudelis, MHS,RN; Elke Kuder-Buletta, RN; Christoph Laske, PhD; Allan Levey, MD, PHD; Johannes Levin, MD; Yan Li, PHD; Oscar Lopez MD, MD; Jacob Marsh, BA; Ralph Martins, PhD; Neal Scott Mason, PhD; Colin Masters, MD; Kwasi Mawuenyega, PhD; Austin McCullough, PhD Candidate; Eric McDade, DO; Arlene Mejia, MD; Estrella Morenas-Rodriguez, MD, PhD; John Morris, MD; James Mountz, MD; Cath Mummery, PhD;N eelesh Nadkarni, MD, PhD; Akemi Nagamatsu, RN; Katie Neimeyer, MS; Yoshiki Niimi, MD;James Noble, MD; Joanne Norton,MSN, RN, PMHCNS-BC ; Brigitte Nuscher, ; Ulricke Obermüller, ; Antoinette O'Connor, MRCPI; Riddhi Patira , MD; Richard Perrin, MD, PhD; Lingyan Ping, PhD; Oliver Preische, MD; Alan Renton, PhD; John Ringman, MD; Stephen Salloway, MD; Peter Schofield, PhD; Michio Senda, MD, PhD; Nicholas T Seyfried, D.Phil; Kristine Shady, BA, BS; Hiroyuki Shimada, MD, PhD; Wendy Sigurdson, RN; Jennifer Smith, PhD; Lori Smith, PA-C; Beth Snitz, PhD; Hamid Sohrabi, PhD; Sochenda Stephens, BS, CCRP; Kevin Taddei, BS ; Sarah Thompson, PA-C; Jonathan Vöglein, MD; Peter Wang, PhD; Qing Wang, PhD; Elise Weamer, MPH; Chengjie Xiong, PhD; Jinbin Xu, PhD; Xiong Xu, BS, MS;
